# Real-time simultaneous monitoring of multiple analytes in bacterial cultures

**DOI:** 10.1101/2025.01.10.632470

**Authors:** Maggie M. Fink, Zahra Aljuboori, Maggie K. Klaers, Morgan Underdue, Joseph Malkovsky, Shahir S. Rizk

**Affiliations:** Department of Biological Sciences, University of Notre Dame, Notre Dame, Indiana, USA; Department of Chemistry and Biochemistry, Indiana University, South Bend, IN, USA; Department of Biochemistry, Molecular Biology and Biophysics, University of Minnesota, Minneapolis, MN, USA; Department of Biochemistry and Molecular Biology, Indiana University School of Medicine, Indianapolis, IN, USA

**Keywords:** Biosensor, Multi-sensor Array, Bacterial Periplasmic Binding Proteins, Bacterial Metabolism, Fluorescence Biosensor

## Abstract

Bacterial metabolites are essential for biological processes, influencing human health, ecosystems, and industrial applications. Simultaneous real-time monitoring of these metabolites is critical in understanding microbial dynamics, particularly in bioreactors and food or drug manufacturing. Current approaches often rely on offline methods, which are labor-intensive and susceptible to contamination, or genetic engineering techniques limited to single-analyte monitoring. Here, we present a novel method utilizing engineered periplasmic binding proteins (PBPs) conjugated with fluorophores to track multiple metabolites simultaneously in *Escherichia coli* cultures. This system continuously monitors the levels of multiple analytes such as glucose, arabinose, ribose, glutamate, and arginine, providing high temporal resolution while maintaining sensor stability over 24 hours. Our findings confirm hierarchical substrate utilization in *E. coli* and demonstrate the versatility of PBP-based multi-sensor arrays. This approach offers a non-invasive, modular, and scalable tool for bacterial metabolite analysis, paving the way for advances in both fundamental discoveries and practical applications in microbiology.

**Importance:** Real-time monitoring of metabolites in bacterial cultures is crucial for advancing our understanding of microbial physiology, metabolic fluxes, and dynamic responses to environmental changes. This capability enables researchers to capture transient metabolic states that are often missed in endpoint measurements. The use of engineered periplasmic binding proteins as biosensors for this real-time metabolite monitoring represents a groundbreaking approach. By leveraging the natural specificity and high affinity of PBPs for small molecules, these biosensors can be engineered to detect a wide range of metabolites with exceptional sensitivity and temporal resolution. The integration of PBP-based biosensors into microbial research not only enhances our ability to study real-time metabolism but also provides a versatile tool for optimizing industrial bioprocesses and exploring bacterial infections and complex microbial ecosystems

## Introduction

Bacterial metabolites are crucial for various biological processes and significantly impact human health and the environment (1–5). These small molecules, produced or consumed during bacterial metabolism, can serve as essential nutrients, signaling molecules, or antimicrobial agents, influencing the microbial community and host physiology (6, 7). In the human body, for example, gut bacteria produce metabolites that regulate immune function, maintain gut integrity, and influence brain health (4, 6–9). Bacterial metabolites also play a vital role in ecosystems, contributing to nutrient cycling and supporting plant growth by promoting root development or protecting against pathogens (8, 10–13). Understanding and harnessing these metabolites can lead to advances in medicine, agriculture, and biotechnology (14).

The ability to identify and simultaneously track multiple metabolites in bacterial cultures is also important in bioreactors used in the food and drug industries (15). The sensors deployed in the complex environment of a growth medium must provide sufficient accuracy, specificity, and sensitivity. Additionally, the sensors must possess the stability to withstand the duration of the experiment or prep while having an appropriate response time. Currently, several types of spectroscopies, electrochemistry, and mass spectrometry are employed as reliable methods to identify a wide range of metabolites (16–23). But often these methods require “off-line” monitoring, where samples are extracted from the culture at different time points for analysis. These “off-line” measurements can be time and labor-intensive and may result in introducing contamination to the cultures (15). While some real-time monitoring of metabolites in bacterial samples has been constructed, they rely on genetic engineering to introduce fluorescence reporters to bacterial strains which are time intensive, not universally applicable, and are often limited to monitoring single metabolites (24–28).

Here, we describe the use of engineered periplasmic binding proteins (PBPs) conjugated with various fluorophores to simultaneously detect multiple metabolites over time in *Escherichia coli* monocultures. PBPs comprise a superfamily of receptor proteins that bind to and recognize a wide variety of metabolites ranging from sugars, amino acids, ions, metals, and even aromatic byproducts of lignin degradation (29, 30). Bacteria species use PBPs to scavenge molecules from the environment using a two-component system that regulates nutrient uptake, gene expression, and chemotaxis (31, 32). A characteristic feature of PBPs is that ligand-binding is associated with a large conformational change through a hinge bending motion from an open unbound to a closed-bound conformation (33, 34). Many PBPs have previously been engineered into biosensors by specific attachment of an environmentally sensitive fluorophore near the hinge region or the binding pocket (29, 35). As a result, the conformational change associated with binding triggers a change in fluorescence indicating ligand concentration.

Our method utilizes PBPs to monitor several analytes in bacterial cultures continuously. We use PBP-fluorophore conjugates to detect single, double, or triple analytes while monitoring bacterial growth. Exogenous addition of the sensor proteins shows the ability to continuously monitor signals over 24 hours in planktonic culture using a fluorescence plate reader, with changes in the signals corresponding to the depletion or production of the tested metabolites, glucose, arabinose, ribose, glutamate, arginine, and ornithine. We show that the engineered biosensors remain stable and sensitive over time, ensuring accurate monitoring of metabolites. Our findings provide a foundation for engineering protein-based, metabolite-specific multi-sensor arrays that can help track and quantify metabolites in complex mixtures without the need for sampling or strain-specific genetic approaches.

## Results

To detect multiple analytes, we characterized the changes in fluorescence associated with ligand binding of several PBP-fluorophore conjugate combinations. We used Glucose binding protein (TnGBP) from *Thermotoga neapolitana*(36), arabinose binding protein (ABP) from *E. coli* (29), ribose binding protein (TteRBP) from *Thermoanaerobacter tengcongensis* (37), and aspartate/glutamate binding protein (EBP) from *E coli*(*29*) to detect changes in glucose, arabinose, ribose, and glutamate concentrations, respectively. A combination of previously published (29) and new cysteine mutations were introduced to the various PBPs for conjugation with fluorophores for the detection of changes in respective ligand binding.

Changes in fluorescence were determined by testing multiple fluorophores to determine the optimal signal range. For tnGBP, conjugation with IANBD showed a large increase in fluorescence emission upon glucose binding (Fig. 1B). Alexa488 and acrylodan gave an increase in fluorescence with ligand binding for ABP (Fig. 1D) and TteRBP (Fig. 1F), respectively. A decrease in fluorescence was observed for EBP conjugated with coumarin when bound to glutamate (Fig. 1H). The dissociation constant (*K*_d_) of each PBP for its ligand was determined by monitoring fluorescence change as a function of ligand concentration (Figure 1). *K*_d_ values ranged between 11µM and 6.4 mM (supplementary information table S1).

**Figure 1:**
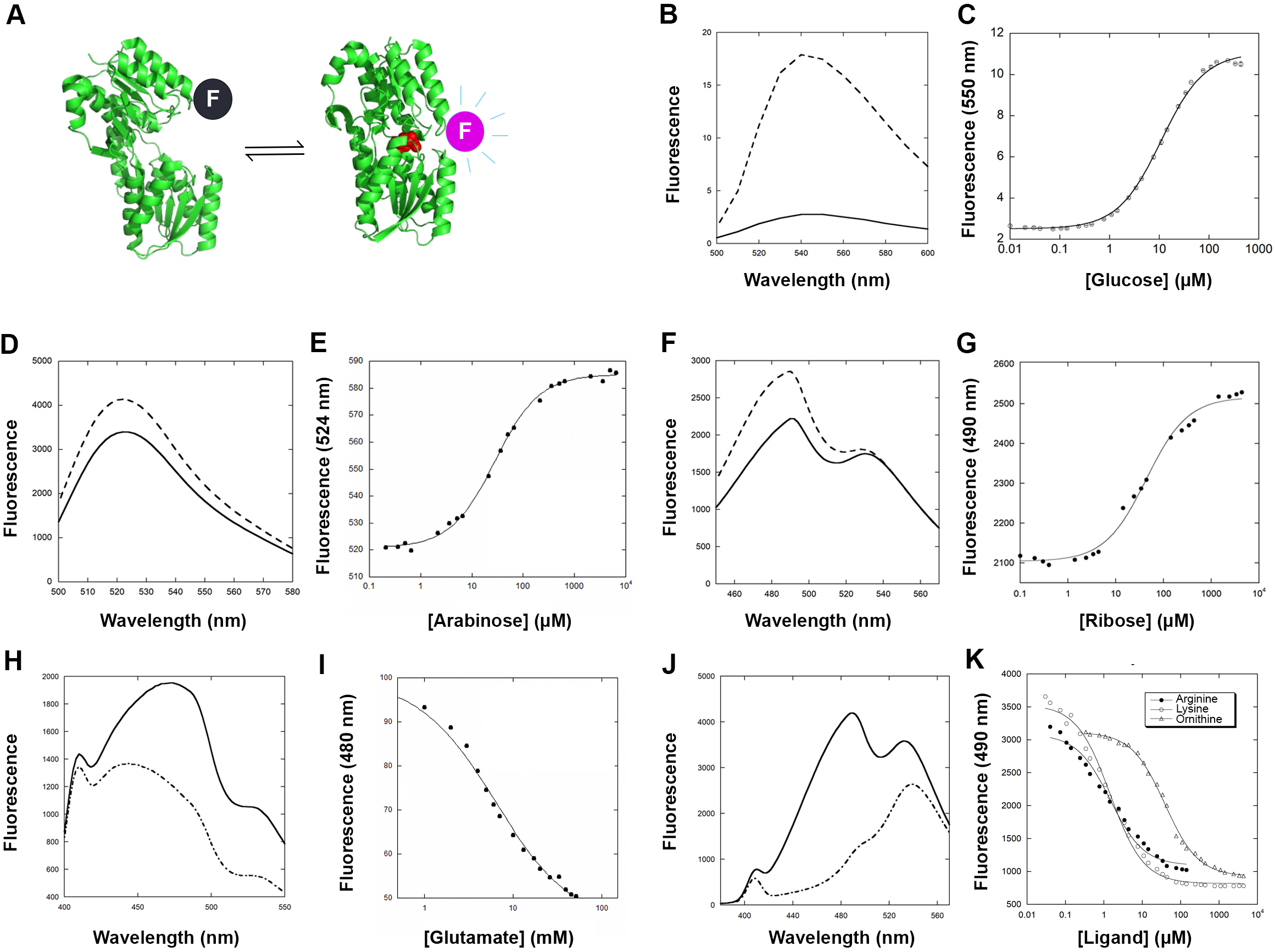
Characterization of PBP-fluorophore conjugates. **A:** PBPs undergo a large hinge-bending conformational change upon binding their ligand (red). The change in fluorescence emission of a conjugated fluorophore (F) can be used as an indication of ligand binding. Shown here as an example is a thermophilic glucose binding protein (PDB codes: 3C6Q and 2H3H). **B—K:** Fluorescence intensity of PBP-fluorophore conjugates in the absence (solid lines) and in the presence of ligand (dashed lines), followed by titration curves. TnGBP-IANDB (B and C), ABP-Alexa488 (D and E), TteRBP-Acrylodan (F and G), EBP-Coumarin (H and I), LAOBP-Acrylodan (J and K).

To develop a biosensor for arginine, we used the lysine/arginine/ornithine binding protein from *Salmonella* (LAOBP) (38). We carried out cysteine scanning by introducing 4 individual cysteine mutations at positions near the hinge guided by the crystal structures of the apo and bound forms of the protein (39) (supplementary information Fig. S1). Each of the 4 mutants was conjugated to 6 different fluorophores, and the fluorescence change upon the addition of ligand was characterized for each of the 24 protein-ligand conjugates. Of the 4 mutants, only the LAOBP-22C mutant showed changes in fluorescence >10% with 4 of the 6 fluorophores tested (Table 1). We determined the dissociation constants for all 3 ligands (arginine, lysine, and ornithine) using the LAOBP-22C acrylodan conjugate (Fig. 1F) and the coumarin conjugate (supplementary information Fig. S2). With a large change in fluorescence and a dissociation constant of 1.5µM for both arginine and lysine, we chose to use the LAOBP-22C acrylodan conjugate for subsequent experiments.

**Table 1:**
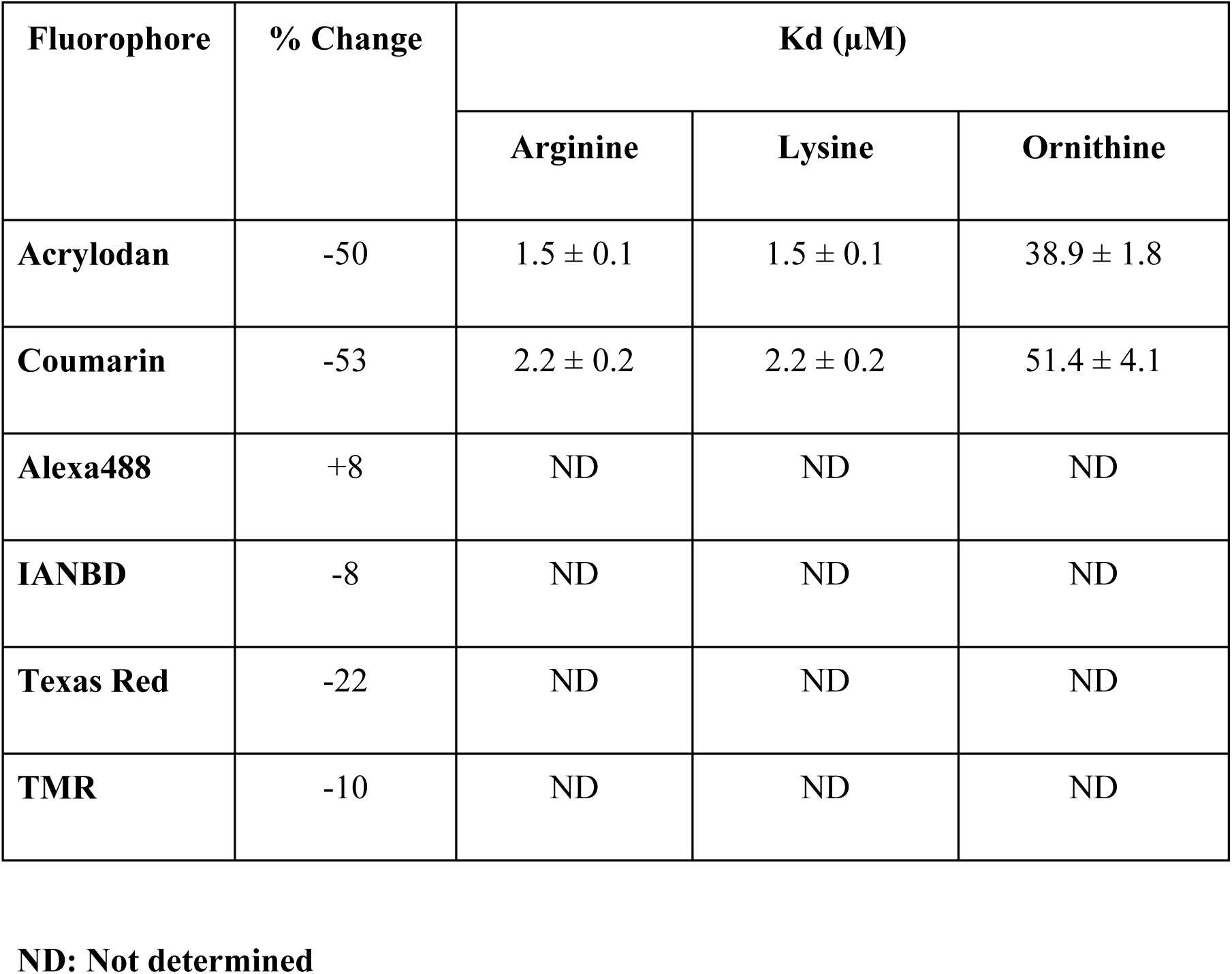
Characterization of the LAOBP-22C mutant conjugated with different fluorophores.

To determine if the PBPs could be used to continuously monitor analytes in complex mixtures over time, 100 nM of fluorescently labeled PBPs were added to *E. coli* K12 cultures grown on minimal media containing defined carbon sources in a 96-well format. In media with glucose (1 mM) and IANBD-labeled TnGBP, fluorescence sharply decreased over a period of 20 minutes corresponding to the mid-log phase of *E. coli* growth when glucose uptake and utilization occurs rapidly(2, 40, 41) (Fig. 2A). During growth on arabinose (1mM), the change in ABP-Alexa488 fluorescence was more gradual, which was consistent with the observed corresponding delay in log phase (Fig. 2B). Cultures grown on ribose (1mM) in the presence of Acrylodan-labeled TteRBP showed a rapid decrease in fluorescence during log phase growth (Fig. 2C). Acrylodan-labeled LAOBP added to media containing only arginine (1mM), which is not a sufficient carbon source for *E. coli*(42), did not exhibit any change in fluorescence or OD (Fig. 2D). However, the use of ornithine (1 mM), which can be utilized to some extend by *E. coli* (43), resulted in a fluorescence change as OD increased at around 18 hours (Fig. 2E). Similarly, in media containing only glutamate (1mM), which is a poor carbon source (43), and EBP-Coumarin, *E. coli* growth was delayed but a change in fluorescence was observed corresponding to an increase in OD (Figure 5F). These results indicate that PBPs can detect the changes in corresponding individual analytes in real-time during *E. coli* growth and reflect current understanding up metabolite uptake and utilization.

**Figure 2.**
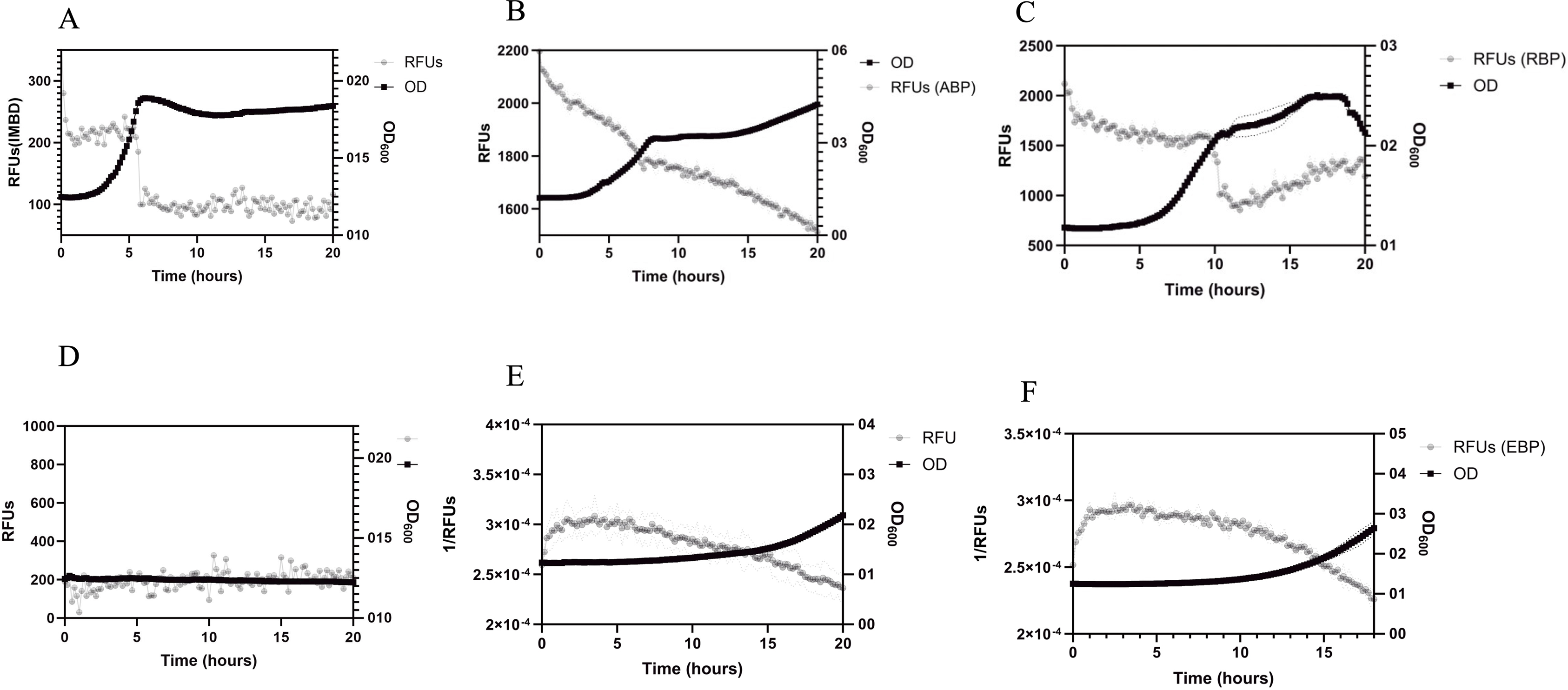
Engineered PBP based biosensors can monitor metabolites in *E. coli* planktonic growth. *E. coli* was grown in 96-well plates in minimal media containing 1 mM of either glucose (A), arabinose (B), ribose, (C), arginine (D), ornithine (E), or glutamate (F). 100 nM of the corresponding labeled PBPs, tnGBP-IANBD, ABP-Alexa488, TteRBP-acrylodan, LAOBP-acrylodan, and EBP-Coumarin, was added to each culture. OD600 and fluorescence was measured every 10 minutes. The change in fluorescence (RFUs; left y-axis) corresponds with *E. coli* growth (OD600; right y-axis) and depletion of each metabolite, except for arginine, which is unable to be utilized as a carbon source.

*E. coli* was also grown in minimal media with different combinations of carbon sources and corresponding PBPs to monitor multiple analytes simultaneously. We grew *E. coli* on glucose and arginine, glucose and ribose, glucose and glutamate, glutamate and arabinose, or glucose, arabinose, and ribose (Fig. 3A-D). For each of the conditions, PBPs were able to detect changes in specific analytes as they were depleted from the media. Our results are consistent with known hierarchical carbon utilization in *E. coli* (44–49). Specifically, in media with glucose and ribose, diauxic growth of *E. coli*(*50*) was observed with changes in fluorescence signals from tnGBP and TteRBP corresponding to two different exponential growth phases (Fig. 3A). In media containing glucose and glutamate, *E. coli* growth was slightly delayed; however, the decrease in tnGBP fluorescence corresponded with log phase, while EBP signal showed a gradual decrease as glutamate was utilized by *E. coli* (51, 52) (Fig. 3B). While LAOBP did not show any change in fluorescence when arginine was the sole carbon source (Fig. 2D), when glucose was added, LAOBP fluorescence change correlated with an increase in *E.coli* growth as glucose was depleted from the media (42) (Fig. 3C). Cultures growing on arabinose and glutamate (Fig. 3D) showed a similar trend to what was observed in media with glucose and glutamate. Both conditions showed a delayed log phase and changes in fluorescence corresponding to *E. coli* growth.

**Figure 3:**
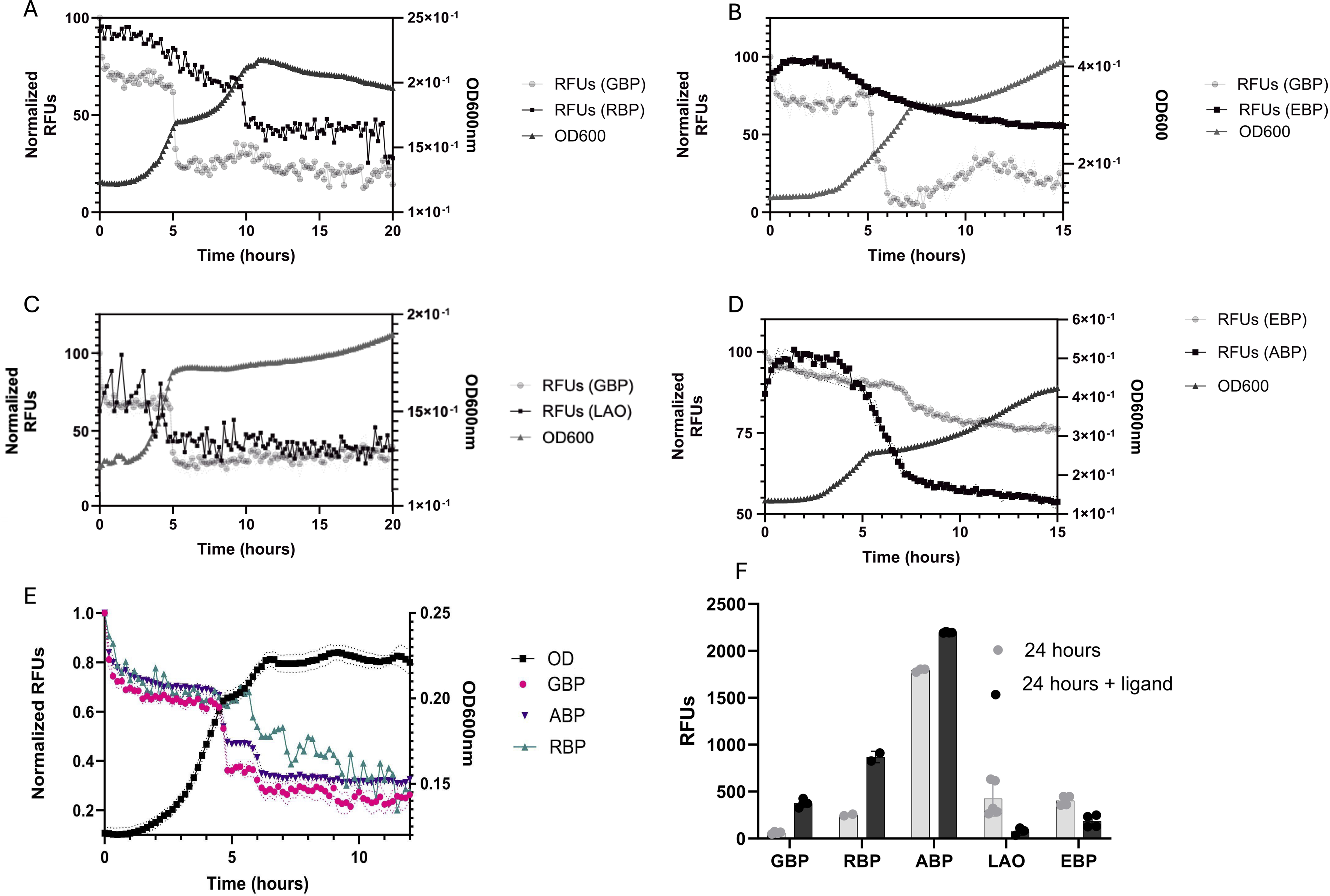
Multiple metabolites can be monitored simultaneously. *E. coli* was grown in minimal media containing 1mM of glucose and glutamate (A), arabinose and glutamate (B), glucose and ribose (C), or glucose and arginine (D), and glucose, arabinose, and ribose (E) and 100 nM of each corresponding labeled PBPs, tnGBP-IANBD, ABP-Alexa488, TteRBP-acrylodan, LAOBP-acrylodan, and EBP-Coumarin Fluorescence for each biosensor was measured every 10 minutes as well as OD600. Changes in fluorescence indicating metabolite consumption are consistent with *E. coli’s* known hierarchical carbon utilization, indicating the metabolite-specific activity of each biosensor. (F) Ligands corresponding to each PBP, glucose, ribose, arabinose, arginine and glutamate were added directly to cultures and fluorescence was measured again. Change in fluorescence (increase or decrease) was consistent Figure 1, indicating that PBPs are still active after 24 hours in *E. coli* culture and that changes in fluorescence are a result of analyte concentration in media.

Finally, *E. coli* was grown in media with three different sugars: glucose, arabinose, and ribose along with their corresponding PBPs (Figure 3E). Decreases in fluorescence were observed for each of the PBPs, again corresponding to *E. coli* diauxic growth phases known to occur on mixed carbon substrates (5, 44–47). The change in fluorescence was also sequential with tnGBP decreasing first, followed by ABP and finally TteRBP which is consistent with previous studies characterizing hierarchical carbon utilization in *E. coli* (44, 46, 47).

To determine if the observed changes in fluorescence were not the result of degradation of the protein sensors over the course of *E. coli* growth, glucose, arabinose, ribose, glutamate, arginine and ornithine were added to 24-hour old cultures containing each corresponding PBP. Immediately after the addition of the respective analyte, fluorescence was measured again (Fig. 3F). Each of the PBPs showed a change in fluorescence, indicating that the sensors are still active, and the changes observed over time in bacterial culture are a result of changes in analytes present in the media.

## Discussion

The method described here represents a non-invasive way to continuously monitor multiple nutrients simultaneously in bacterial cultures. By employing the exquisite specificity of PBPs and their characteristic conformation change upon ligand binding, each PBP acts as an independent sensor in a multisensor array, providing real-time information on how each nutrient is consumed. Other methods, like mass spectrometry, require the collection of time points, which can be labor-intensive and can potentially introduce contamination. Our method bypasses the need for collecting time points allowing high temporal resolution within minutes over hours or days.

The use of PBPs as the biosensor elements in the multisensor array described here offers several advantages. First, the superfamily of PBPs offers a wide repertoire of ligand specificity, making this method modular and expandable to include a wide variety of molecules not only those consumed by bacteria, but others that are produced such as those that serve as signaling molecules. With binding kinetic constants in the sub-millisecond time-scale (53), PBPs can quickly respond to a change in ligand concentration providing high temporal resolution. Second, careful cysteine scanning provides multiple potential positions within each protein where one of several fluorophores can be attached, each showing a significant change in fluorescence upon binding to the ligand. This allows a mix-and-match approach, where different PBPs can be combined in several ways to detect different combinations of ligands with each PBP acting as a unique channel. Third, we show here that the PBP-fluorescent conjugates are stable and retain their ligand-binding capabilities for the entire duration of the experiment. Utilizing thermophilic PBPs offers an additional advantage for use under harsh conditions where other protein-based biosensors may become denatured. Finally, PBPs are highly engineerable; mutations near and within the binding pocket can be introduced to fine-tune affinity of the protein for its ligand, allowing a tunable range for ligand detection (54). Furthermore, adding conformation-specific antibodies can also modulate the affinity of PBPs by controlling the equilibrium of the conformational change (55, 56). In some cases, PBPs have been modified to bind to different ligands (57) or entirely re-engineered to bind synthetic molecules (58), further expanding their application in real-time sensing.

We also demonstrated the ability to detect the sequential consumption of 3 different sugars while monitoring the growth of *E. coli* using a fluorescent plate reader to measure fluorescence and OD_600_. Our data shows with high temporal resolution that when presented with three different sugars, glucose is consumed first, followed by arabinose, then ribose, consistent with known regulation mechanisms of *E. coli’s* sequential utilization of carbon sources. Using the method described here can contribute significantly to understanding bacterial metabolism and can help decipher the role of different metabolites in bacterial growth, communication, and pathogenesis (12, 59, 60).

Our results also provide the potential for studying bacterial metabolism beyond monocultures. By adapting this approach, it can be applied to multispecies cultures with the goal of understanding the temporal dynamics of interspecies bacterial communication mediated through chemical signals (61, 62). This has far-reaching implications in understanding infections, biofilm formation, the mechanisms of antibiotic resistance, and how commensal species compete or cooperate within host environments (63–66). Furthermore, with some modifications, this method could be extended to study nutrient uptake and metabolite release in a wide range of infection models where altered metabolic pathways create unique nutrient environments that impact infection progression and severity.

## Acknowledgements

This work was supported by the Research Corporation for Science Advancement Cottrell Scholars Award #25865 (to SR); MF was funded by the NSF GRFP, ZA and MU were funded by the Carolyn and Lawrence Garber Summer Research Scholarship; JM was supported by the Louis Stokes Alliances for Minority Participation program through the National Science Foundation Grant EES 2308500. We thank Prof. Joshua Shrout for access to instrumentation.

## Materials and Methods

### Cloning and site-directed mutagenesis

The gene coding for TnGBP was cloned from *Thermotoga neapolitana* genomic DNA (ATCC) by PCR using primers that introduce an N-terminal NdeI site and a C-terminal NheI site. Genes for LAOBP, EBP, TtRBP, and ABP were ordered from IDT. PCR was used to amplify the genes using universal primers that introduce an N-terminal NdeI site and C-terminal NheI site. All genes were cloned into pET25b+ cut with NdeI and NheI-HF in-frame with the plasmid sequence which introduces a C-terminal HSV tag followed by a 6-His tag. Cloning was confirmed by DNA sequencing.

### Protein expression and purification

Plasmids coding for each of the proteins were used to transform BL21-DE3 *E. coli* cells by heat shock (45 sec, at 42°C), and plated on ampicillin LB agar plates. Single colonies were used to inoculate overnight cultures of 2XYT-amp, which were used to inoculate 250 mL 2XYT-amp in 1 L baffled flasks. After reaching OD600 of ∼0.6-0.8, protein expression was induced by IPTG (1mM final concentrations) and cultures were allowed to grow for an additional 3-4 hrs at 37°C or overnight at 16°C. Cells were isolated by centrifugation, resuspended in 20 mM TRIS-HCl, 500 mM NaCl, 10 mM imidazole pH 8.6 (buffer A), lysed by addition of lysozyme and sonication in the presence of DNAse, followed by centrifugation at 8,000 rpm for 30 min. The cleared lysate was loaded on a 1 or 5 mL His-Trap Nickel-charged column (Cytiva) equilibrated with buffer A on an ÄKTA start. Proteins were isolated using a 25 mL gradient between buffer A and buffer B (20 mM TRIS-HCl, 500 mM NaCl, 10 mM imidazole pH 8.6). SDS-PAGE analysis was used to identify fractions containing pure protein. Econo-Pac 10DG columns (Bio-Rad) were used to exchange the protein into working buffer (50 mM MOPS, 150 mM NaCl, pH 6.9).

### Fluorophore conjugation

A 5-10-fold excess of fluorophore to protein ratio was used to attach thiol-reactive fluorophores (dissolved in DMSO) to the cysteine residue of each of the PBPs. Reaction was carried out at room temperature overnight in the presence of TCEP (1 mM final concentration) tumbling end to end. Excess fluorophore was removed using Econo-Pac 10DG columns equilibrated with working buffer.

### Fluorescent screening and data analysis

Fluorescence analysis of protein-fluorophore conjugates was carried out on a Jasco FP-8500 fluorometer. UV-clear 4-sided cuvettes containing 200-300 nM labeled protein in working buffer were used. The following parameters were used for each of the fluorophores: Acrylodan (excitation: 359 nm, emission: 440-550 nm), Coumarin (excitation: 390 nm, emission: 430-520 nm), IANBD (excitation: 460 nm, emission: 500-600 nm), Alexa488 (excitation: 490 nm, emission: 500-560 nm), TMR (excitation: 555 nm, emission: 570-620 nm), Texas Red (excitation: 595 nm, emission: 600-660 nm).

Titrations were carried out by adding increasing amounts of ligand while monitoring changes in emission at the maximum wavelength. Fluorescence values were volume adjusted, and Kd values were determined by fitting the titration data using Kaleidagraph according to the following equation:

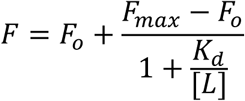

where, where *F* is the observed fluorescence at a given ligand concentration [L], *F_o_*is the fluorescence in the absence of ligand, and *F_max_* is the fluorescence at saturation.

### Bacterial strains and culture conditions

*E. coli* strains used in this study are included in Table S1. Cultures were streaked from frozen (−80°C) stocks onto lysogeny broth (LB) agar plates (1.0% wt/vol) and incubated at 37°C overnight. Isolated colonies were selected and inoculated in a 6 mL LB. Tubes were incubated overnight at 37°C with shaking at 240 rpm. Cells were centrifuged at 10,000 rpm for 20 minutes and washed with 6 mL PBS and washed a second time before resuspension in 6 mL M9 with 1mM carbon source. LB broth or Minimal Media (M9) was used unless otherwise described.

### Growth Curves

Growth curves were generated in clear 96 well plates (Corning Incorporated). And incubated at 37°C in a PowerWave_X_ 340 (BIO-TEK INSTRUMENTS, INC) Cultures were inoculated into 1 mL of Mueller-Hinton broth (Sigma-Aldrich) or FAB to an OD_600_ of 0.01. 200 uL/well was added to 96-well black clear bottom plates (Corning Incorporated) for each culture condition and absorbance at 600 nm was measured every 10 minutes for 24 hours at 37°C. The plate was shaken for 10 seconds before each read. 100 nM of biosensor was added to each culture condition. Relative fluorescent intensity was captured at excitation (add in wavelengths) every ten minutes for 24 hours at 37°C. The plate was shaken for 10 seconds before each read.

## References

1. Eisenreich W, Dandekar T, Heesemann J, Goebel W. 2010. Carbon metabolism of intracellular bacterial pathogens and possible links to virulence. Nat Rev Microbiol 8:401–12.

2. Gorke B, Stulke J. 2008. Carbon catabolite repression in bacteria: many ways to make the most out of nutrients. Nat Rev Microbiol 6:613–24.

3. Moreno R, Rojo F. 2023. The importance of understanding the regulation of bacterial metabolism. Environ Microbiol 25:54–58.

4. Postler TS, Ghosh S. 2017. Understanding the Holobiont: How Microbial Metabolites Affect Human Health and Shape the Immune System. Cell Metab 26:110–130.

5. Wedmark YK, Vik JO, Oyas O. 2024. A hierarchy of metabolite exchanges in metabolic models of microbial species and communities. PLoS Comput Biol 20:e1012472.

6. Hosseinkhani F, Heinken A, Thiele I, Lindenburg PW, Harms AC, Hankemeier T. 2021. The contribution of gut bacterial metabolites in the human immune signaling pathway of non-communicable diseases. Gut Microbes 13:1–22.

7. Krautkramer KA, Fan J, Backhed F. 2021. Gut microbial metabolites as multi-kingdom intermediates. Nat Rev Microbiol 19:77–94.

8. Franzino T, Boubakri H, Cernava T, Abrouk D, Achouak W, Reverchon S, Nasser W, Haichar FEZ. 2022. Implications of carbon catabolite repression for plant-microbe interactions. Plant Commun 3:100272.

9. Haran JP, Bhattarai SK, Foley SE, Dutta P, Ward DV, Bucci V, McCormick BA. 2019. Alzheimer’s Disease Microbiome Is Associated with Dysregulation of the Anti-Inflammatory P-Glycoprotein Pathway. mBio 10.

10. Bulgarelli D, Schlaeppi K, Spaepen S, Ver Loren van Themaat E, Schulze-Lefert P. 2013. Structure and functions of the bacterial microbiota of plants. Annu Rev Plant Biol 64:807–38.

11. Pacheco AR, Vorholt JA. 2023. Resolving metabolic interaction mechanisms in plant microbiomes. Curr Opin Microbiol 74:102317.

12. Passalacqua KD, Charbonneau ME, O’Riordan MXD. 2016. Bacterial Metabolism Shapes the Host-Pathogen Interface. Microbiol Spectr 4.

13. Xian L, Yu G, Wei Y, Rufian JS, Li Y, Zhuang H, Xue H, Morcillo RJL, Macho AP. 2020. A Bacterial Effector Protein Hijacks Plant Metabolism to Support Pathogen Nutrition. Cell Host Microbe 28:548–557 e7.

14. DeBerardinis RJ, Keshari KR. 2022. Metabolic analysis as a driver for discovery, diagnosis, and therapy. Cell 185:2678–2689.

15. Abdelfattah A, Hossain MI, Cheng L. 2020. High-strength wastewater treatment using microbial biofilm reactor: a critical review. World J Microbiol Biotechnol 36:75.

16. Meier S, Jensen PR, Duus JO. 2011. Real-time detection of central carbon metabolism in living Escherichia coli and its response to perturbations. FEBS Lett 585:3133–8.

17. Miyawaki A, Llopis J, Heim R, McCaffery JM, Adams JA, Ikura M, Tsien RY. 1997. Fluorescent indicators for Ca2+ based on green fluorescent proteins and calmodulin. Nature 388:882–7.

18. Qiu S, Cai Y, Yao H, Lin C, Xie Y, Tang S, Zhang A. 2023. Small molecule metabolites: discovery of biomarkers and therapeutic targets. Signal Transduct Target Ther 8:132.

19. Lai Z, Tsugawa H, Wohlgemuth G, Mehta S, Mueller M, Zheng Y, Ogiwara A, Meissen J, Showalter M, Takeuchi K, Kind T, Beal P, Arita M, Fiehn O. 2018. Identifying metabolites by integrating metabolome databases with mass spectrometry cheminformatics. Nat Methods 15:53–56.

20. Bolten CJ, Kiefer P, Letisse F, Portais JC, Wittmann C. 2007. Sampling for metabolome analysis of microorganisms. Anal Chem 79:3843–9.

21. Bolten CJ, Wittmann C. 2008. Appropriate sampling for intracellular amino acid analysis in five phylogenetically different yeasts. Biotechnol Lett 30:1993–2000.

22. Bagnara AS, Finch LR. 1972. Quantitative extraction and estimation of intracellular nucleoside triphosphates of Escherichia coli. Anal Biochem 45:24–34.

23. Imperlini E, Santorelli L, Orru S, Scolamiero E, Ruoppolo M, Caterino M. 2016. Mass Spectrometry-Based Metabolomic and Proteomic Strategies in Organic Acidemias. Biomed Res Int 2016:9210408.

24. Koveal D, Rosen PC, Meyer DJ, Diaz-Garcia CM, Wang Y, Cai LH, Chou PJ, Weitz DA, Yellen G. 2022. A high-throughput multiparameter screen for accelerated development and optimization of soluble genetically encoded fluorescent biosensors. Nat Commun 13:2919.

25. Ovechkina VS, Zakian SM, Medvedev SP, Valetdinova KR. 2021. Genetically Encoded Fluorescent Biosensors for Biomedical Applications. Biomedicines 9.

26. Mo GC, Ross B, Hertel F, Manna P, Yang X, Greenwald E, Booth C, Plummer AM, Tenner B, Chen Z, Wang Y, Kennedy EJ, Cole PA, Fleming KG, Palmer A, Jimenez R, Xiao J, Dedecker P, Zhang J. 2017. Genetically encoded biosensors for visualizing live-cell biochemical activity at super-resolution. Nat Methods 14:427–434.

27. Palmer AE, Qin Y, Park JG, McCombs JE. 2011. Design and application of genetically encoded biosensors. Trends Biotechnol 29:144–52.

28. Greenwald EC, Mehta S, Zhang J. 2018. Genetically Encoded Fluorescent Biosensors Illuminate the Spatiotemporal Regulation of Signaling Networks. Chem Rev 118:11707–11794.

29. de Lorimier RM, Smith JJ, Dwyer MA, Looger LL, Sali KM, Paavola CD, Rizk SS, Sadigov S, Conrad DW, Loew L, Hellinga HW. 2002. Construction of a fluorescent biosensor family. Protein Sci 11:2655–75.

30. Bisson C, Salmon RC, West L, Rafferty JB, Hitchcock A, Thomas GH, Kelly DJ. 2022. The structural basis for high-affinity uptake of lignin-derived aromatic compounds by proteobacterial TRAP transporters. FEBS J 289:436–456.

31. Borrok MJ, Zhu Y, Forest KT, Kiessling LL. 2009. Structure-based design of a periplasmic binding protein antagonist that prevents domain closure. ACS Chem Biol 4:447–56.

32. Groisman EA. 2016. Feedback Control of Two-Component Regulatory Systems. Annu Rev Microbiol 70:103–24.

33. Bermejo GA, Strub MP, Ho C, Tjandra N. 2010. Ligand-free open-closed transitions of periplasmic binding proteins: the case of glutamine-binding protein. Biochemistry 49:1893–902.

34. Kroger P, Shanmugaratnam S, Ferruz N, Schweimer K, Hocker B. 2021. A comprehensive binding study illustrates ligand recognition in the periplasmic binding protein PotF. Structure 29:433–443 e4.

35. Rizk SS, Cuneo MJ, Hellinga HW. 2006. Identification of cognate ligands for the Escherichia coli phnD protein product and engineering of a reagentless fluorescent biosensor for phosphonates. Protein Sci 15:1745–51.

36. Tian Y, Cuneo MJ, Changela A, Hocker B, Beese LS, Hellinga HW. 2007. Structure-based design of robust glucose biosensors using a Thermotoga maritima periplasmic glucose-binding protein. Protein Sci 16:2240–50.

37. de Lorimier RM, Tian Y, Hellinga HW. 2006. Binding and signaling of surface-immobilized reagentless fluorescent biosensors derived from periplasmic binding proteins. Protein Sci 15:1936–44.

38. Nikaido K, Ames GF. 1992. Purification and characterization of the periplasmic lysine-, arginine-, ornithine-binding protein (LAO) from Salmonella typhimurium. J Biol Chem 267:20706–12.

39. Kang CH, Shin WC, Yamagata Y, Gokcen S, Ames GF, Kim SH. 1991. Crystal structure of the lysine-, arginine-, ornithine-binding protein (LAO) from Salmonella typhimurium at 2.7-A resolution. J Biol Chem 266:23893–9.

40. Carreon-Rodriguez OE, Gosset G, Escalante A, Bolivar F. 2023. Glucose Transport in Escherichia coli: From Basics to Transport Engineering. Microorganisms 11.

41. Negrete A, Ng WI, Shiloach J. 2010. Glucose uptake regulation in E. coli by the small RNA SgrS: comparative analysis of E. coli K-12 (JM109 and MG1655) and E. coli B (BL21). Microb Cell Fact 9:75.

42. Schneider BL, Kiupakis AK, Reitzer LJ. 1998. Arginine catabolism and the arginine succinyltransferase pathway in Escherichia coli. J Bacteriol 180:4278–86.

43. Neidhardt FC, Curtiss R. 1996. Escherichia coli and Salmonella: cellular and molecular biology, 2nd ed. ASM Press, Washington, D.C.

44. Okano H, Hermsen R, Kochanowski K, Hwa T. 2020. Regulation underlying hierarchical and simultaneous utilization of carbon substrates by flux sensors in Escherichia coli. Nat Microbiol 5:206–215.

45. Okano H, Hermsen R, Hwa T. 2021. Hierarchical and simultaneous utilization of carbon substrates: mechanistic insights, physiological roles, and ecological consequences. Curr Opin Microbiol 63:172–178.

46. Aidelberg G, Towbin BD, Rothschild D, Dekel E, Bren A, Alon U. 2014. Hierarchy of non-glucose sugars in Escherichia coli. BMC Syst Biol 8:133.

47. Ammar EM, Wang X, Rao CV. 2018. Regulation of metabolism in Escherichia coli during growth on mixtures of the non-glucose sugars: arabinose, lactose, and xylose. Sci Rep 8:609.

48. Deutscher J. 2008. The mechanisms of carbon catabolite repression in bacteria. Curr Opin Microbiol 11:87–93.

49. Kremling A, Geiselmann J, Ropers D, de Jong H. 2015. Understanding carbon catabolite repression in Escherichia coli using quantitative models. Trends Microbiol 23:99–109.

50. Salvy P, Hatzimanikatis V. 2021. Emergence of diauxie as an optimal growth strategy under resource allocation constraints in cellular metabolism. Proc Natl Acad Sci U S A 118.

51. Bren A, Park JO, Towbin BD, Dekel E, Rabinowitz JD, Alon U. 2016. Glucose becomes one of the worst carbon sources for E.coli on poor nitrogen sources due to suboptimal levels of cAMP. Sci Rep 6:24834.

52. Reitzer L. 2003. Nitrogen assimilation and global regulation in Escherichia coli. Annu Rev Microbiol 57:155–76.

53. Miller DM, 3rd, Olson JS, Pflugrath JW, Quiocho FA. 1983. Rates of ligand binding to periplasmic proteins involved in bacterial transport and chemotaxis. J Biol Chem 258:13665-72.

54. Marvin JS, Hellinga HW. 2001. Manipulation of ligand binding affinity by exploitation of conformational coupling. Nat Struct Biol 8:795–8.

55. Rizk SS, Paduch M, Heithaus JH, Duguid EM, Sandstrom A, Kossiakoff AA. 2011. Allosteric control of ligand-binding affinity using engineered conformation-specific effector proteins. Nat Struct Mol Biol 18:437–42.

56. Mukherjee S, Griffin DH, Horn JR, Rizk SS, Nocula-Lugowska M, Malmqvist M, Kim SS, Kossiakoff AA. 2018. Engineered synthetic antibodies as probes to quantify the energetic contributions of ligand binding to conformational changes in proteins. J Biol Chem 293:2815–2828.

57. Banda-Vazquez J, Shanmugaratnam S, Rodriguez-Sotres R, Torres-Larios A, Hocker B, Sosa-Peinado A. 2018. Redesign of LAOBP to bind novel l-amino acid ligands. Protein Sci 27:957–968.

58. N’Guetta PY, Fink MM, Rizk SS. 2020. Engineering a fluorescence biosensor for the herbicide glyphosate. Protein Eng Des Sel 33.

59. Nogales J, Garmendia J. 2022. Bacterial metabolism and pathogenesis intimate intertwining: time for metabolic modelling to come into action. Microb Biotechnol 15:95–102.

60. King AN, de Mets F, Brinsmade SR. 2020. Who’s in control? Regulation of metabolism and pathogenesis in space and time. Curr Opin Microbiol 55:88–96.

61. Shetty SA, Kuipers B, Atashgahi S, Aalvink S, Smidt H, de Vos WM. 2022. Inter-species Metabolic Interactions in an In-vitro Minimal Human Gut Microbiome of Core Bacteria. NPJ Biofilms Microbiomes 8:21.

62. Culp EJ, Goodman AL. 2023. Cross-feeding in the gut microbiome: Ecology and mechanisms. Cell Host Microbe 31:485–499.

63. Zhang Y, Cai Y, Zhang B, Zhang YPJ. 2024. Spatially structured exchange of metabolites enhances bacterial survival and resilience in biofilms. Nat Commun 15:7575.

64. Yeor-Davidi E, Zverzhinetsky M, Krivitsky V, Patolsky F. 2020. Real-time monitoring of bacterial biofilms metabolic activity by a redox-reactive nanosensors array. J Nanobiotechnology 18:81.

65. Stokes JM, Lopatkin AJ, Lobritz MA, Collins JJ. 2019. Bacterial Metabolism and Antibiotic Efficacy. Cell Metab 30:251–259.

66. Baquero F, Martinez JL. 2017. Interventions on Metabolism: Making Antibiotic-Susceptible Bacteria. mBio 8.

